# Stimulating human prefrontal cortex increases reward learning

**DOI:** 10.1101/2021.01.27.428488

**Authors:** Margot Juliëtte Overman, Verena Sarrazin, Michael Browning, Jacinta O’Shea

**Author notes:** Contributed equally to this paper (Michael Browning, Jacinta O’Shea). Corresponding author: Jacinta O’Shea, University of Oxford, Warneford Hospital, Oxford OX3 7JX, England.

## Abstract

Work in computational psychiatry suggests that mood disorders may stem from aberrant reinforcement learning processes. Specifically, it is proposed that depressed individuals believe that negative events are more informative than positive events, resulting in faster learning from negative outcomes (Pulcu & Browning, 2019). In this proof-of-concept study, we investigated whether learning rates for affective outcomes are malleable using transcranial direct current stimulation (tDCS). Healthy adults completed an established reinforcement learning task (Pulcu & Browning, 2017) in which the information content of reward and loss outcomes was manipulated by varying the volatility of stimulus-outcome associations. Learning rates on the tasks were quantified using computational models. Stimulation over dorsolateral prefrontal cortex (DLPFC) but not motor cortex (M1) specifically increased learning rates for reward outcomes. The effects of prefrontal tDCS were cognitive state-dependent: online stimulation increased learning rates for wins; offline stimulation decreased both win and loss learning rates. A replication study confirmed the key finding that online tDCS to DLPFC specifically increased learning rates for rewards relative to losses. Taken together, these findings demonstrate the potential of tDCS for modulating computational parameters of reinforcement learning relevant to mood disorders.

**Significance statement:** Disproportionate learning from negative relative to positive outcomes has been implicated in the development and maintenance of depression. The present work demonstrates that transcranial direct current stimulation (tDCS) to dorsolateral prefrontal cortex can specifically increase learning from positive events in healthy adults. Our results provide preliminary evidence that non-invasive brain stimulation can be used to shape reinforcement learning, indicating a potential novel cognitive neurostimulation intervention strategy for affective disorders.

## Introduction

The ability to learn from our experiences is central to adaptive decision making. Computational accounts of reinforcement learning posit that optimal learners should determine which events are most *informative* and weight these accordingly when making choices (MacKay, 2003; Behrens et al., 2007; Nassar et al., 2012; Browning et al., 2015). The information content of an event depends in part on the volatility of the association being learned. When action-outcome contingencies change frequently, each new observation is relatively informative about the current state of the association. Predictions should therefore be updated more rapidly for volatile than stable associations (Pulcu & Browning, 2019). In keeping with this theory, healthy adults flexibly adapt their learning rates to match the volatility of the environment (Behrens et al., 2007; Nassar et al., 2012; Browning et al., 2015). Moreover, humans can maintain separate estimates for the information content of positive and negative events (Pulcu & Browning, 2017).

An emerging line of computational research suggests that aberrant tracking of these statistical properties could form a core mechanism underpinning affective disorders (Browning et al., 2015; Pulcu & Browning, 2019). Building on preclinical work, it is proposed that depression may stem from a tendency to overestimate the information content of negative relative to positive events (Pulcu & Browning, 2019). This could in turn lead to a negative cognitive bias, which has been causally linked to depression (Mathews & MacLeod, 2005; Eshel & Roiser, 2010). Behaviourally, distorted estimates of information content may reduce an individual’s ability to select actions associated with beneficial outcomes, thereby creating progressively poorer environments. As an illustrative example, a belief that negative events are highly informative may cause an individual to pay more attention to criticism than praise at work. As well as impairing mood and self-confidence, this belief is likely to increase the influence of negative feedback on a person’s decision to pursue or give up on a potentially fruitful project. From this perspective, rebalancing learning from rewarding versus aversive outcomes could be a promising novel approach to ameliorating negative cognitive biases that causally maintain symptoms of depression.

Depression is consistently linked with hypoactivity of the left dorsolateral prefrontal cortex (DLPFC) and concurrent hyperactivity of the right DLPFC (Grimm et al., 2008; Koenigs & Grafman, 2009; Disner et al., 2011). DLPFC also forms part of distributed reward learning circuitry (Haber & Knutson, 2010; Lee et al., 2012) with recent reports pointing to a role in tracking the volatility of reward outcomes (Massi et al., 2018; Farahashi et al., 2019). Hence, modulating activity in DLPFC could influence both the neural and cognitive mechanisms underlying depression. Previously, clinical trials have demonstrated that transcranial direct current stimulation (tDCS) over bilateral DLPFC induces positive but small effects on depressive symptoms (Brunoni et al., 2016; Shiozawa et al., 2014). In these trials, neurostimulation is typically administered while the patient is at rest. However, tDCS applied *during learning* has been shown to strengthen memory for what is being learned - thought to be caused by stabilization of synapses undergoing activity-dependent long-term potentiation (Reis et al., 2009; Fritsch et al., 2010; O’Shea et al., 2017). Hence, the functional impact (and therapeutic potential) of tDCS could be enhanced by stimulating while simultaneously engaging and reshaping learning processes that underpin negative affective biases.

Here we performed a proof-of-concept test of this hypothesis via a series of experiments in healthy adults. We predicted that tDCS over DLPFC would increase reward learning rates compared with sham tDCS. We further predicted that this effect would be cognitive-state specific, induced by tDCS during learning but not by tDCS applied before learning. Finally, we hypothesized that tDCS effects would be anatomically specific, with stimulation to prefrontal but not motor cortex (M1) selectively increasing reward learning rates. To measure reward and loss learning rates we used an established reinforcement learning paradigm in which volatility of stimulus-outcome associations is varied across blocks (Browning et al., 2015; Pulcu & Browning, 2017; Pulcu et al., 2019). This task enabled us to test for potential valence- and volatility-specific effects of stimulation. We applied tDCS bilaterally over DLPFC with a single dose of the stimulation montage and protocol commonly used in depression treatment trials.

## Methods and Materials

### Participants

Eighty healthy native English-speaking adults (45 female, mean age = 24.71, SD ± 5.08) were recruited from the community via local advertisements for four independent studies. Exclusion criteria were left-handedness, a history of psychiatric disorders, neurological illness, use of psychoactive medication, personal or family history of epileptic fits or seizures, and any other contraindications to tDCS. The experimental protocol was approved by the University of Oxford Central University Ethics Committee (RE48995/RE002) and all participants gave written informed consent prior to the study. Demographic details of the participants are provided in Table 1.

**Table 1.**
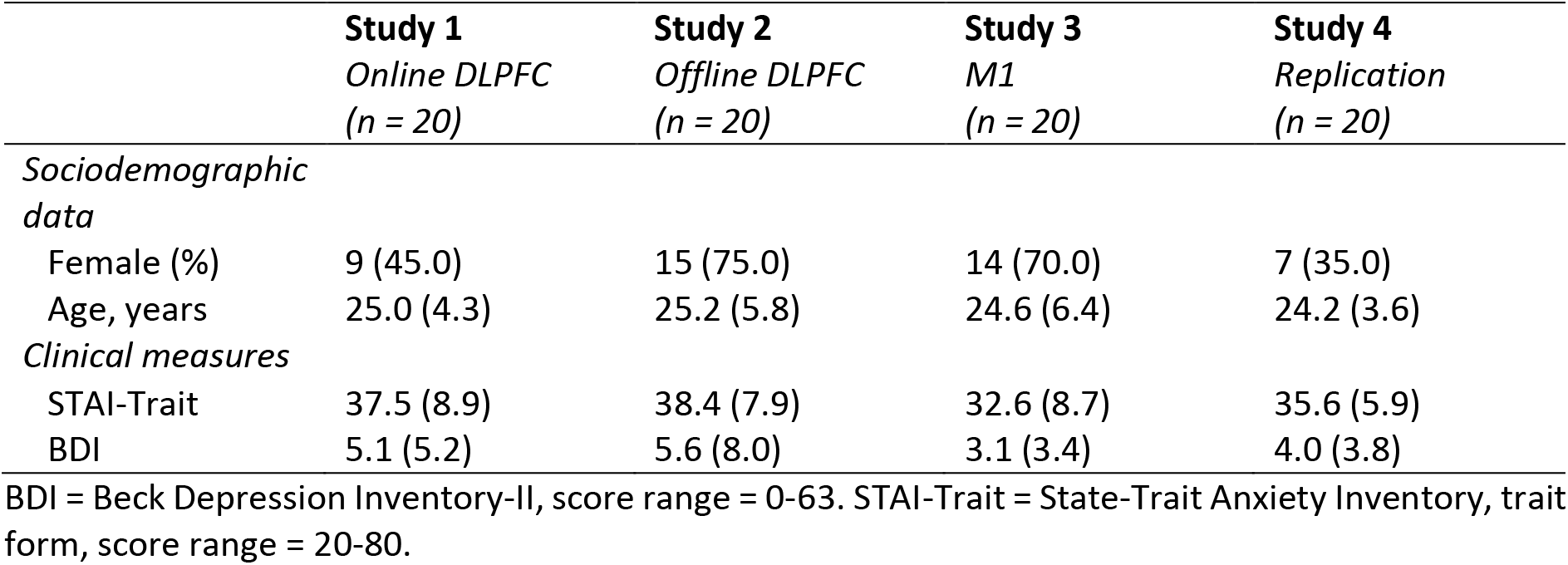
Mean (SD) baseline characteristics by tDCS group.

|  | **Study 1**<br>*Online DLPFC*<br>(*n* = 20) | **Study 2**<br>*Offline DLPFC*<br>(*n* = 20) | **Study 3**<br>*M1*<br>(*n* = 20) | **Study 4**<br>*Replication*<br>(*n* = 20) |
| --- | --- | --- | --- | --- |
| *Sociodemographic data* |  |  |  |  |
| Female (%) | 9 (45.0) | 15 (75.0) | 14 (70.0) | 7 (35.0) |
| Age, years | 25.0 (4.3) | 25.2 (5.8) | 24.6 (6.4) | 24.2 (3.6) |
| *Clinical measures* |  |  |  |  |
| STAI-Trait | 37.5 (8.9) | 38.4 (7.9) | 32.6 (8.7) | 35.6 (5.9) |
| BDI | 5.1 (5.2) | 5.6 (8.0) | 3.1 (3.4) | 4.0 (3.8) |

BDI = Beck Depression Inventory-II, score range = 0-63. STAI-Trait = State-Trait Anxiety Inventory, trait form, score range = 20-80.

### Study overview and experimental design

Participants took part in one of four stimulation studies, each consisting of two tDCS and reward learning sessions. All participants underwent both active and sham tDCS sessions in a cross-over, double-blind design. Stimulation order was counterbalanced in all groups and sessions were scheduled at least one week apart to minimise carryover effects of repeated learning and/or tDCS. In Study 1, tDCS was applied to DLPFC during task performance to test for the predicted increase in reward learning. In Study 2, tDCS was applied to DLPFC prior to task performance to determine the cognitive state dependence of the DLPFC stimulation effect. In Study 3, online tDCS was delivered over primary motor cortex (M1) to assess the anatomical specificity of the stimulation effects. Study 4 aimed to replicate the findings of Study 1 to evaluate the consistency of behavioural changes induced by online prefrontal tDCS.

### Questionnaires

Symptoms of depression and anxiety (see Table 1) were assessed at baseline with the Beck Depression Inventory-II (BDI) (Beck et al., 1996) and the Trait subscale of the State-Trait Anxiety Inventory (STAI-Trait) (Spielberger et al., 1983). Higher scores on these tasks indicate greater symptoms of depression and anxiety, respectively. For the BDI, cut-off scores suggested by Beck and colleagues (1996) are 0-13 for no or minimal depression, 14-19 for mild depression, 20-80 for moderate depression, and 29-63 for severe depression. The STAI-Trait was not designed for clinical diagnosis, and therefore has no formal cut-off points. To monitor potential changes in acute mood and anxiety across the tDCS sessions, participants completed the Positive and Negative Affect Scales (PANAS) (Watson et al., 1988) and the State-Trait Anxiety Inventory (STAI-State) (Spielberger et al., 1983) immediately before and after completion of the cognitive task (see Information Bias Learning Task (IBLT) below). All scores and analyses of the PANAS and STAI-State are reported in the Extended Data.

### Information Bias Learning Task (IBLT)

The Information Bias Learning Task (IBLT) is a computerised reinforcement learning paradigm which has been described in detail previously (Browning et al., 2015; Pulcu & Browning, 2017; Pulcu et al., 2019). The IBLT was presented on a laptop computer using Presentation® software (Neurobehavioral Systems, Inc., Berkeley, CA, www.neurobs.com). On every trial, a fixation cross in the centre of the screen was flanked by two abstract shapes (letters selected from the Agathodaimon font). Participants were asked to choose the shape they believed would result in the best outcome via a button press, after which a win (+10p) and a loss (−10p) outcome appeared in randomised order above or below the shapes. Participants’ accumulated total winnings were displayed under the fixation cross and updated at the beginning of the subsequent trial. The win and loss outcomes were independent, such that a specific shape could be associated with one, both, or neither of the outcomes. Participants therefore had to form separate predictions for the likelihood of the win and the loss appearing over a specific shape, and select the optimal choice based on those estimates.

Participants completed six task blocks of 80 trials each, with a fixed 30-second rest period between blocks. The same two shapes were used within a task block, and different shapes were used across task blocks. The volatility of shape-outcome contingencies was varied across task blocks to manipulate the relative information content of the win and loss outcomes (see Figure 1). During volatile (i.e., informative) task periods, the association of an outcome with shape ‘A’ reversed between 20% and 80% every 14 to 30 trials. During stable (i.e., uninformative) task periods, the association of an outcome with shape ‘A’ remained constant at 50%. The probability of an outcome appearing over shape ‘B’ is calculated as 1 – shape ‘A’. In blocks 1 and 6, both the win and loss outcome associations were volatile. The aim of these ‘Both-volatile’ blocks was to measure the extent to which participants preferentially learned from equally informative positive and negative outcomes. In blocks 2-5, one outcome was volatile whereas the other remained stable. These blocks, in which only win or only loss outcomes were highly volatile, will be referred to as ‘Win-volatile’ and ‘Loss-volatile’ blocks, respectively. The Win- and Loss-volatile blocks enabled us to simultaneously test for potential specific effects of tDCS on learning for outcomes of different valences (positive vs. negative) and levels of information content (informative vs. uninformative). Win- and Loss-volatile blocks were alternated, with order of presentation (i.e., Win-volatile or Loss-volatile block first) being counterbalanced across participants in each group (see Figure 2.A).

**Figure 1.**
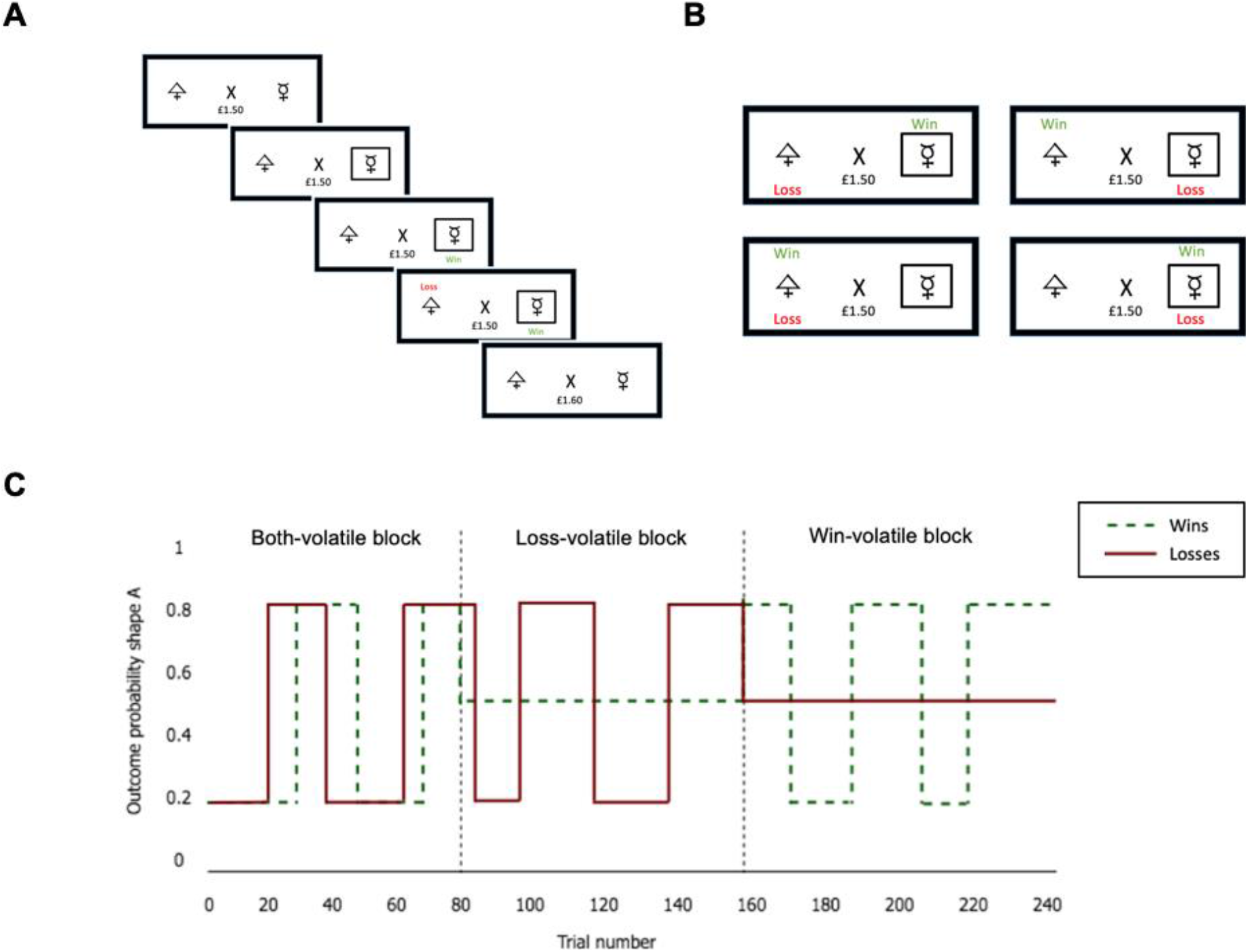
**(A)** Schematic representation of a trial on the Information Bias Learning Task (IBLT; Pulcu & Browning, 2017). After showing the fixation cross and total amount of money won, two abstract shapes are presented on either side of the cross. Once the participant has chosen one of the shapes via a button press, a black frame appears around that shape and a win and loss outcome appear successively in randomised order. A win outcome leads to an increase of 10p, whereas a loss outcome represents a decrease of 10p from the total amount of money won. The total amount of money is updated at the start of the next trial. The aim of the task is to maximise earnings by learning the probabilities of the win and the loss appearing over the respective shapes. **(B)** The four possible outcomes on a task trial. The win and the loss outcomes are independently associated with one of the shapes, allowing for a shape to be associated with one, both, or neither of the outcomes at a given time. **(C)** Volatility of the win (green) and loss (red) outcomes across blocks of the Information Bias Learning Task (IBLT). Volatility for the two outcomes is manipulated independently across task blocks, either switching between 20% and 80% choice-outcome associations or remaining stable at 50% choice-outcome associations. If both wins and losses are volatile, participants should rapidly update their predictions for both types of outcomes (i.e., have a high learning rate). In the ‘Win-volatile’ blocks participants should adopt a high learning rate for wins and a low learning rate for losses, whereas the opposite approach should be taken in the ‘Loss-volatile’ blocks.

**Figure 2.**
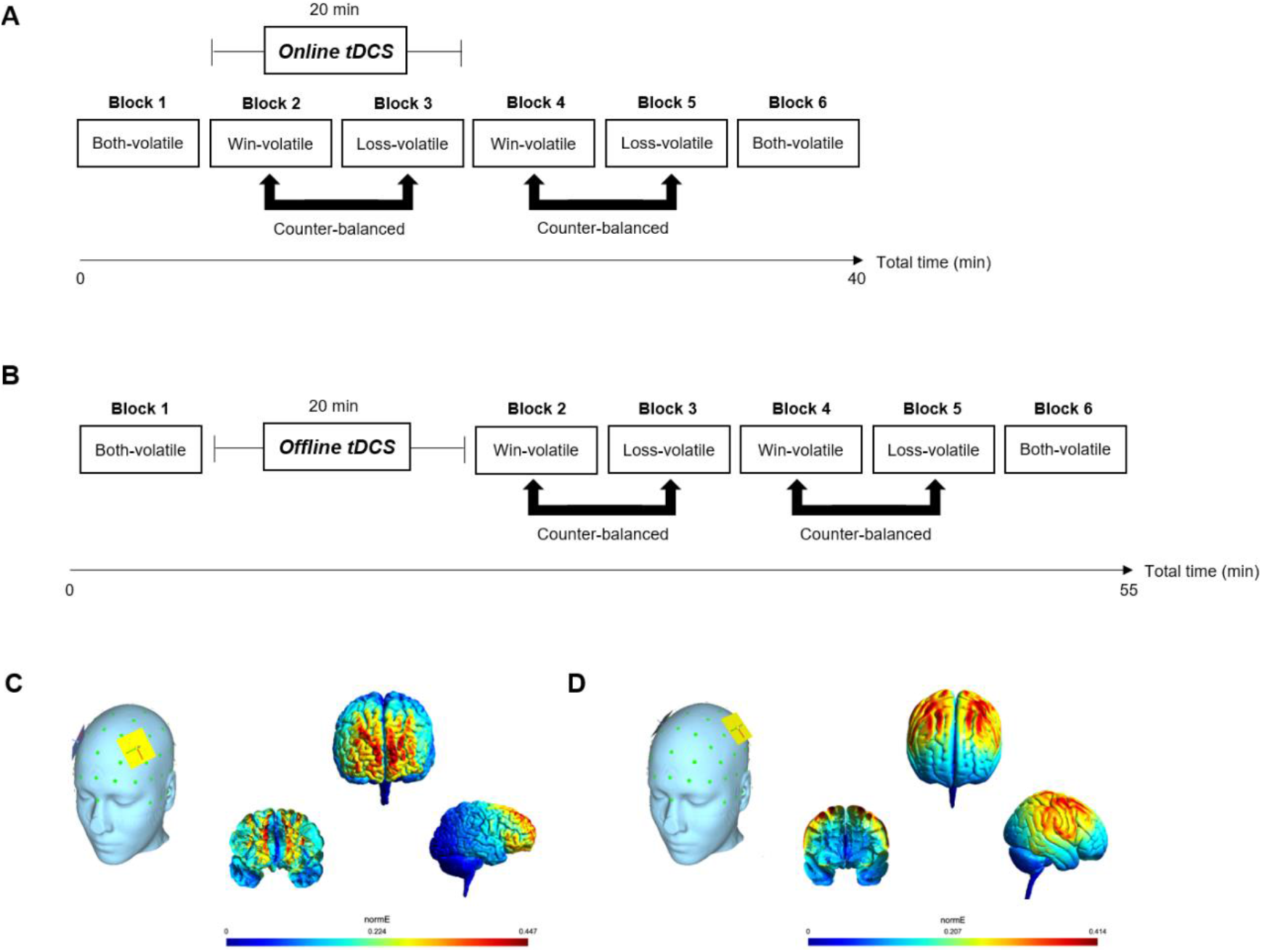
**(A)** Structure and timeline of the online tDCS paradigm (Study 1, 3, and 4). Participants complete six blocks of the IBLT. The task starts and ends with a ‘Both-volatile’ block in which win and loss outcomes are equally informative. The participants are then presented with two ‘Win-volatile’ and two ‘Loss-volatile’ blocks, with block type order counterbalanced across participants. tDCS was applied during Blocks 2-3 of the IBLT. **(B)** Study 2 structure and timeline, in which DLPFC tDCS was applied offline prior to Blocks 2-6 of the Information Bias Learning Task (IBLT). **(C)** Simulation of the electric field induced in the brain by the bilateral prefrontal tDCS montage, with the anodal electrode (yellow) over the left DLPFC (F3) and the cathode (blue) over the right DLPFC (F4). **(D)** Simulation of the electric field induced in the brain by the bilateral motor cortex tDCS montage, with the anodal electrode (yellow) over left M1 and the cathode (blue) over right M1.

In all tDCS studies, block 1 was completed prior to stimulation to provide a baseline measure of learning rates for win and loss outcomes. As win and loss outcomes were equally informative in this block, the learning rates provide an indication of potential learning bias prior to stimulation. In the online tDCS studies (Study 1, 3, and 4), stimulation was applied during IBLT blocks 2 and 3, with blocks 4-6 carried out after tDCS (see Figure 2.A). Blocks 4 and 5 served to test for potential sustained effects post-stimulation. Block 6 (‘Both-volatile’) was used to measure potential changes in learning bias by the end of the task compared with at baseline in Block 1. In the offline tDCS study (Study 2), tDCS was applied first, while participants sat at rest, followed by task blocks 2-6 (see Figure 2.B). The goal of the study was to test if prefrontal stimulation during the task specifically increased learning from positive outcomes. This would manifest as a selective increase in learning rates for wins.

### tDCS protocol and current distribution

Stimulation was delivered using a battery-powered device (Eldith DC-Stimulator-Plus, Neuroconn, Germany). Two rubber electrodes (5 × 5 cm) were placed in saline-soaked sponges and attached to the scalp using rubber bands. For prefrontal stimulation the anodal electrode was placed over the left DLPFC while the cathodal electrode was placed over the right DLPFC (F3 and F4, respectively, according to the 10/20 system of electrode placement). For bilateral stimulation of M1, the anode was centred over the hand area of left primary motor cortex, 5 cm lateral to the vertex, and the cathode over the homologous region of the right hemisphere. In the active tDCS conditions, stimulation was delivered at 2 mA for 20 minutes, with 10s ramping-up and ramping-down. In the sham tDCS conditions, participants received 30s of direct current followed by impedance control with a small current pulse being produced every 550 ms (110 μA over 15 ms), resulting in an instantaneous current of no more than 2 μA. Double-blinding was implemented through the use of a study mode on the tDCS device. SimNIBS (Thielscher et al., 2015) and Gmsh software (Geuzaine & Remacle, 2009) were used to visualize the spatial distribution of the electrical field induced in the brain by the DLPFC and M1 tDCS electrode montages (see Figure 2 C. and 2.D).

### Computational modelling

In line with previous studies utilising the IBLT (Pulcu et al., 2019) we analysed choice behaviour with a model in which a Rescorla-Wagner learning rule (Rescorla & Wagner, 1972) was coupled to a softmax function. The model calculates the probability estimates for the win (*rwin*) and loss (*rloss*) outcomes being associated with shape ‘A’ on the next trial (*i*+1):

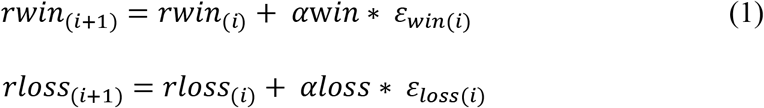

in which *awin* and *aloss* representing the learning rates (value between 0 and 1), and *ε*_*win*(*i*)_ and *ε*_*loss*(*i*)_ represent the prediction error for the win and loss outcomes on the *i*^th^ trial, respectively.
*rwin* and *rloss* were initialised at 0.5 at the start of each block. Estimated outcome probabilities were then transformed into a single choice probability:

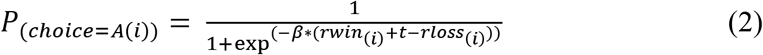

Here, *P*_(*choice*_=_*A*(_*i*_))_ is the probability of choosing shape ‘A’ in trial *i*. *β*represents the inverse decision temperature, or the degree to which the expected values are used to determine choice for a particular shape. Finally, *t* reflects a potential bias towards one of the options over the other. Learning rates and *β*-values were estimated separately for each task block and participant. This was achieved by calculating the full joint posterior probability of the parameters given participants’ choices, deriving the expected value of each parameter from their marginalised probability distributions. The first 10 trials of each block were omitted when fitting the model parameters to participants’ choices, as initial learning rates without prior knowledge were expected to differ from informed learning rates in later trials of the task. The choice of this model was based on a formal comparison using Bayesian information criterion (BIC) values for five competitor models (see Extended Data). Models were implemented using MATLAB version R2018a (The MathWorks, Inc., Natick, MA).

### Non-computational choice behaviour

To test for behavioural effects of tDCS without reliance on the specific assumptions of the model described above, we also conducted non-computational analyses of choice behaviour on the IBLT. In these analyses we focused specifically on choices following trials where one shape was associated with <u>both</u> a win and a loss outcome. On such trials there is no change in earnings because the win and loss cancel each other out. Hence, the shape selected on the next trial provides a measure of the *relative* influence of positive versus negative outcomes on participants’ subsequent choice behaviour. If the positive outcome more strongly influences a participant’s choices, they would be expected to stay with the shape currently associated with both outcomes and choose it again on the next trial. By contrast, if the negative outcome is more influential, the participant would be expected to switch on the next trial and instead choose the other shape that was not associated with both outcomes. The proportion of trials in which the participants chose the win-over the loss-driven option was calculated for each block. Trials in which the win and loss outcome were associated with different shapes were excluded from the non-model-based analyses, as in these trials both outcomes promote the same choice (i.e. selecting the shape associated with the win outcome).

### Statistical analyses

All analyses were completed in R software (Version 3.6.0). Learning rates and inverse temperature values derived from the computational model, non-computational choice behaviour, and total winnings (in £) were entered into repeated-measures ANOVAs with the ‘ezANOVA’ function from the ‘ez’ R package. The first set of analyses assessed effects of online prefrontal tDCS on reward learning (Study 1). Next, we carried out control comparisons to confirm whether tDCS outcomes were timing- and site-specific. Here, Cognitive State (online vs. offline, i.e. Study 1 vs. Study 2) or tDCS Target (DLPFC vs. M1, i.e. Study 1 vs. Study 3) was included as a between-subjects variable. Finally, data from Study 4 (replication study) were analysed in a separate ANOVA to establish whether effects of online prefrontal stimulation were robust.

As in previous work (Pulcu et al., 2019), outcome measures from the Win- and Loss-volatile blocks of the IBLT were the primary focus of analysis. The main outcome of interest was participants’ learning behaviour in the Win- and Loss-volatile blocks, which were tested for outcome- and volatility-dependent effects of stimulation. The independent variables were tDCS, Outcome, Time (during/post-tDCS = blocks 2-3/4-5), and Volatility (Win-volatile/Loss-volatile blocks). The dependent variable was learning rate.

Both-volatile blocks were analysed to test for potential changes in learning from the baseline pre-tDCS Block (1) to the final block post-tDCS/task (6). The independent variables were tDCS (active/sham), Outcome (win/loss), and Time (block 1/block 6). Baseline learning rates for wins and losses from Block 1 of the first of the two test sessions were included as a covariate, to account for individual differences in baseline learning rate biases.

Inverse temperature values, non-model-based choice behaviour, and total winnings were also investigated with repeated-measures ANOVAs, with the independent variables of tDCS, Time, and Volatility. Stimulation order (sham or active tDCS first) was included as a between-subjects variable in all analyses. The effect sizes for all ANOVAs are reported as generalised eta squared values (*η*^2^G). Significant interactions were followed up with *post hoc* paired *t*-tests. Learning rates were transformed onto the infinite real line using an inverse logistic transform, and inverse temperature values were normalised with a log transformation consistent with previous studies (Pulcu & Browning, 2017; Pulcu et al., 2019). Figures and reported values represent raw parameter values to facilitate interpretation of the results.

### Data availability statement

Raw data, analysis scripts, and task materials can be accessed on Open Science Framework: https://osf.io/av9pf/, DOI: 10.17605/OSF.IO/AV9PF.

## Results

### Online prefrontal tDCS increases reward learning rates

#### Computational parameters

We predicted that online stimulation of DLPFC would selectively enhance reward learning. In line with this hypothesis, we observed a valence-specific effect of prefrontal tDCS in Win- and Loss-volatile blocks (tDCS × Outcome interaction: *F*_(1,18)_ = 4.89, *p* = 0.040; *η*^2^G = 0.006) (Figure 3.A). Active tDCS caused higher learning rates for win (*t*_(19)_ = 2.11, *p* = 0.048) but not loss outcomes (*t*_(19)_ = 0.35, *p* = 0.728). This effect did not change over time (tDCS × Outcome × Time interaction: *F*_(1,18)_ = 0.56, *p* = 0.464), indicating that the increase in reward learning rates was maintained for at least 15 minutes after tDCS (see Extended Data for visualization of learning rates over time). Stimulation effects varied by volatility of the outcomes (tDCS × Volatility interaction: *F*_(1,18)_ = 8.09, *p* = 0.011, *η*^2^G = 0.014), such that tDCS increased learning rates in Loss-volatile (*t*_(19)_ = 2.47, *p* = 0.023) but not Win-volatile blocks (*t*_(19)_ = −0.12, *p* = 0.905). As shown in Figure 4, this effect appears to be most pronounced for win outcomes. Prefrontal stimulation specifically altered learning rates, without changing the randomness of participants’ choices (no effect of tDCS on inverse temperature parameters; all *p* >0.05). For the Both-volatile blocks (Block 1 vs 6), there were no differences in learning rates or inverse temperature values between the active vs sham tDCS sessions (all *p* > 0.05).

**Figure 3.**
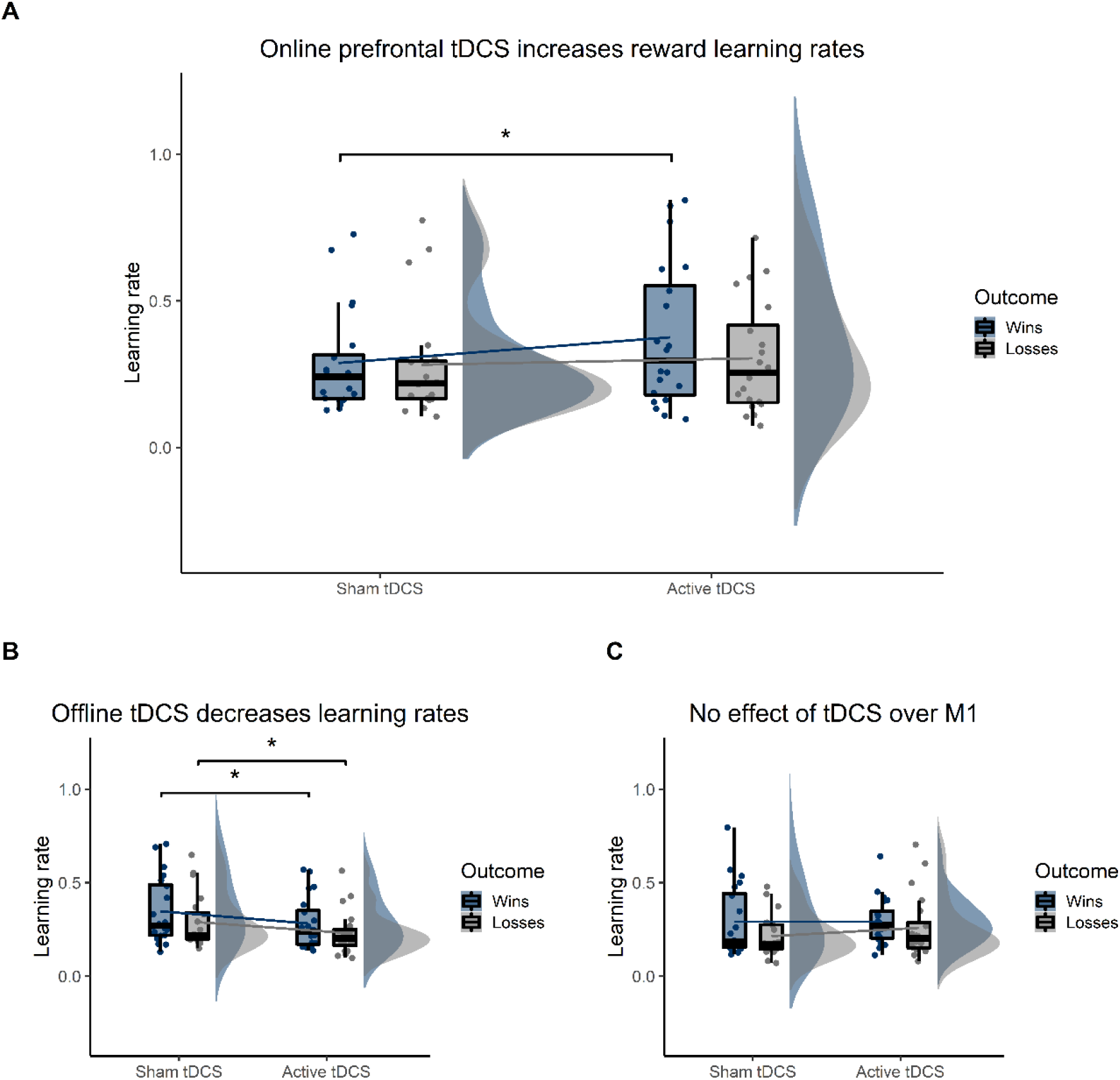
Valence-specific tDCS effects in Win- and Loss-volatile blocks. **(A)** Prefrontal stimulation *during* task performance selectively increased learning rates for win outcomes (\**p* <0.05). **(B)** Effects of prefrontal tDCS were cognitive-state specific, with stimulation applied *before* task performance decreasing both win and loss learning rates (\**p* <0.05). **(C)** tDCS effects on reward learning were anatomically specific, with no effects of stimulation over motor cortex. Violin-plots show the distribution of learning rates by outcome (wins = blue, losses = grey). Summary statistics are provided in boxplots, with the black horizontal line indicating the median and whiskers representing the 25^th^ and 75^th^ percentiles of values. Dots represent participants’ individual data points averaged across task blocks.

**Figure 4.**
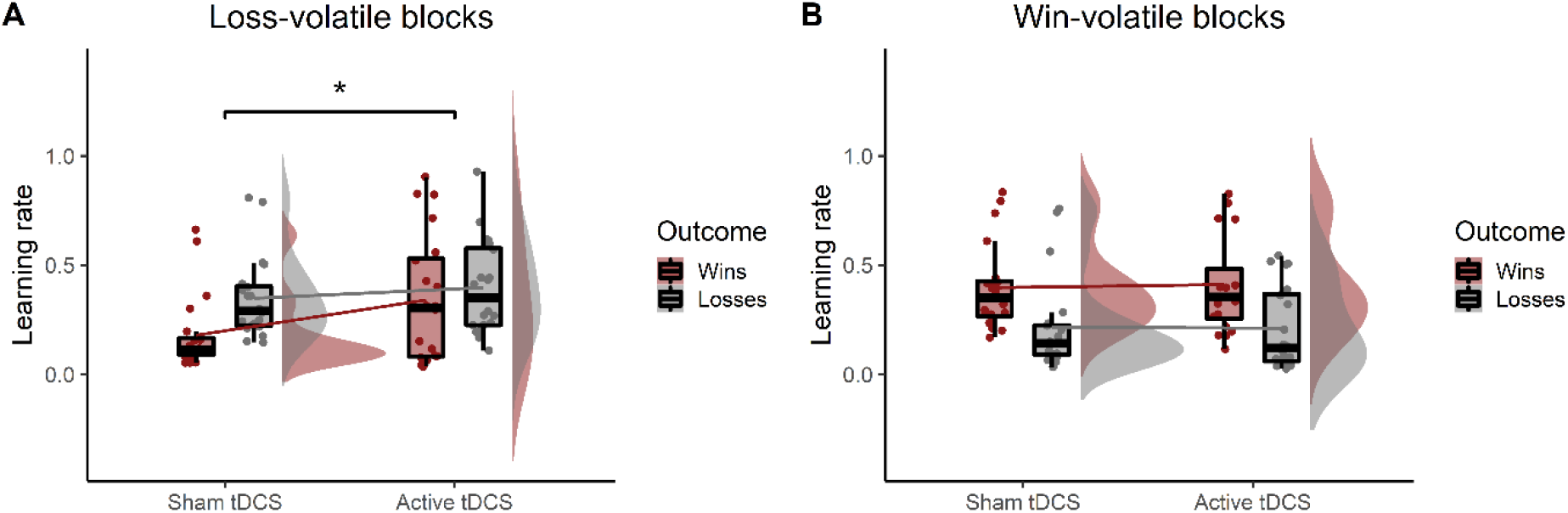
Volatility-dependent effects of online prefrontal tDCS. **(A)** tDCS increased learning rates in Loss-volatile blocks (\**p* <0.05). **(B)** There was no change in learning rates in Win-volatile blocks with prefrontal tDCS. Violin-plots show the distribution of learning rates by outcome type (wins = red, losses = grey). Summary statistics are provided in boxplots, with the black horizontal line indicating the median and whiskers representing the 25^th^ and 75^th^ percentiles of values. Dots represent participants’ individual data points averaged across task blocks.

#### Non-computational choice behaviour and total winnings

For win-driven choices, there was an interaction of tDCS and volatility (*F*_(1,18)_ = 7.90, *p* = 0.016, *η*^2^G = 0.029). Active stimulation increased the number of win-driven choices in Loss-volatile blocks (*t*_(19)_ = 3.33, *p* = 0.004), but had no effect in Win-volatile blocks (*t*_(19)_ = −0.71, *p*= 0.489). Thus, in line with the computational modelling results, tDCS to DLPFC increased learning from reward outcomes, with effects particularly prominent when win outcomes were less informative than loss outcomes (i.e. in the Loss-volatile blocks). For total winnings, we found a main effect of tDCS (*F*_(1,18)_ = 5.91, *p* = 0.026, *η*^2^G = 0.023). Overall, participants won less money with active than sham stimulation. This decrease in total winnings is consistent with participants’ increasing reliance on uninformative rewards. In the Both-volatile blocks, there were no effects of tDCS (all *p* >0.05).

### Opposing effects of online and offline stimulation

To test for cognitive state specificity, in Study 2 we applied tDCS offline prior to task performance while participants simply rested and compared the effects of online versus offline stimulation. Crucially, online and offline tDCS had diverging effects on win and loss learning rates (Cognitive State × tDCS × Outcome interaction: *F*_(1,36)_ = 5.10, *p* = 0.030, *η*^2^G = 0.003). In contrast with the specific increase in reward learning with online tDCS, offline tDCS induced a *reduction* of both win and loss learning rates (*t*_(19)_ = −2.18, *p* = 0.042) (Figure 3.B). Contrary to online stimulation, offline tDCS did not affect any of the non-computational outcomes (all *p* >0.05). As predicted, the specific effects of prefrontal tDCS on reward learning were induced only by *online* stimulation.

### Anatomical specificity of tDCS effects

To determine the anatomical specificity of tDCS effects, we compared learning rates with online DLPFC versus M1 stimulation. Importantly, stimulation effects differed by outcome for the two tDCS targets in Win- and Loss-volatile blocks (tDCS Target × tDCS condition × Outcome interaction: *F*_(1,36)_ = 4.29, *p* = 0.045, *η*^2^G = 0.005) (Figure 3.C). Whereas prefrontal stimulation specifically increased overall win learning rates, tDCS over motor cortex had no overall effect on learning rates for either wins (*t*_(19)_ = 0.22, *p* = 0.828) or losses (*t*_(19)_ = 1.47, *p*= 0.159). Thus, the valence-specific increase in reward learning observed with prefrontal stimulation was anatomically specific. In addition, tDCS effects varied depending on the volatility of the outcomes (tDCS target × tDCS condition × Volatility interaction: *F*_(1,36)_ = 13.58, *p* <0.001, *η*^2^G = 0.012). Whereas prefrontal tDCS increased learning rates in the Loss-volatile blocks, M1 stimulation increased learning rates in the Win-volatile blocks (*t*_(19)_ = 2.40, *p* = 0.027). As expected, tDCS over M1 did not alter any of the non-computational outcomes (all *p* >0.05).

### Replication of online tDCS effects

In Study 4, we carried out a replication of Study 1 to determine whether online prefrontal tDCS consistently increases reward learning rates. We did not find the expected outcome valence- or volatility-specific effects of tDCS observed in Study 1 (tDCS × Outcome interaction: *F*_(1,18)_ = 1.30, *p* = 0.269; tDCS × Volatility interaction: *F*_(1,18)_ = 0.01, *p* = 0.930). There was also no significant interaction of tDCS condition with volatility in the non-computational analyses (*F*_(1,18)_ = 0433, *p* = 0.519). However, planned comparisons demonstrated that active tDCS caused higher learning rates for win than loss outcomes in the Win- and Loss-volatile blocks (*t*_(79)_ = 2.18, *p* = 0.032) (Figure 5). This suggests that active prefrontal stimulation was associated with increased learning from positive outcomes compared to sham tDCS, although the effect was weaker in the replication dataset than in the original Study 1.

**Figure 5.**
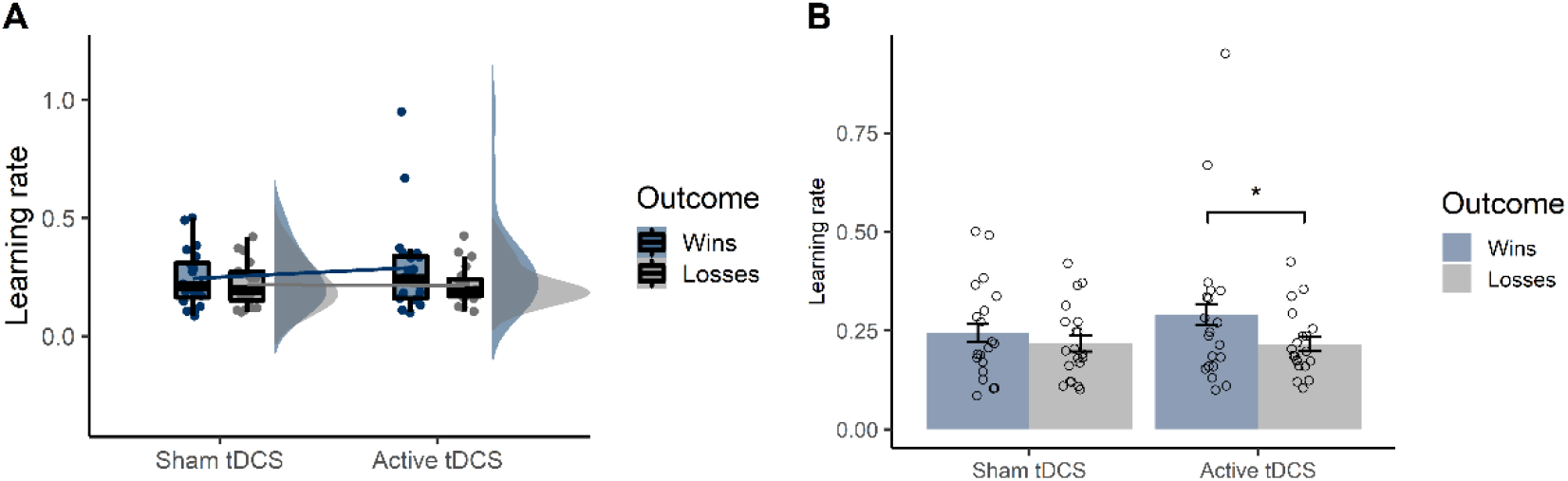
Replication study (Study 4). **(A)** Increased reward learning rates with online prefrontal tDCS were not significant in a repeated-measures ANOVA (tDCS × Outcome interaction: *p* = 0.269). **(B)** Planned comparisons revealed that tDCS caused higher learning rates for wins than losses (*p* <0.05).

## Discussion

This study tested the malleability of reward learning with transcranial direct current stimulation in healthy adults. Online prefrontal tDCS selectively increased reward learning rates, with behavioural changes outlasting stimulation for at least 15 minutes. A replication of this study demonstrated a weaker but similar effect of prefrontal tDCS on reward learning rates. As predicted, tDCS effects were cognitive-state dependent, with offline stimulation resulting in *decreased* learning rates for both wins and losses. In addition, tDCS outcomes were anatomically specific, with no effects of motor cortex stimulation. Taken together, these findings demonstrate the potential of online prefrontal tDCS for shifting the balance of learning from positive and negative outcomes.

### Valence-specific tDCS effects

Our finding of a selective increase in learning from positive outcomes is consistent with several prior stimulation studies targeting left DLPFC, which have reported particular improvements in cognitive control for positive stimuli (Vanderhasselt et al., 2013) and recognition of positive emotions (Nitsche et al., 2012). According to the asymmetry hypothesis of valence, responses to positive and negative stimuli are lateralised to left and right DLPFC, respectively (Davidson, 1992; Herrington et al., 2005; Herrington et al., 2010; Wyczesany et al., 2018). Anodal stimulation of left DLPFC (and concurrent cathodal stimulation of right DLPFC) may therefore predominantly enhance processing of positive information.

It is worth noting that electric field simulations suggested that our tDCS montage may also reach medial regions of prefrontal cortex. Previous studies have linked the medial prefrontal cortex (mPFC) with various aspects of reward learning, including action-outcome predictions (Alexander & Brown, 2011), social prediction errors (Behrens et al., 2008), and belief updating (Kuzmanovic et al., 2018). Changes in learning rates with tDCS could thus be mediated by neural activations in the mPFC. Concurrent tDCS-fMRI studies will be essential to establish the neural mechanisms underpinning the effects of prefrontal stimulation on reward learning.

Alternatively, selective changes in reward learning rates could stem from pre-existing patterns of neural activation and behaviour. Healthy adults tend to have an ‘optimism bias’ (Sharot, 2011) and are more likely to attend to positive compared with negative or neural stimuli (Kress et al., 2016). It is therefore possible that a prior tendency towards learning from positive outcomes was amplified with stimulation in healthy volunteers. A critical next step will be to explore tDCS effects in participants presenting with a range of baseline learning rates.

### Learning from uninformative outcomes

A key feature of the Information Bias Learning Task is that it can measure participants’ responses to informative (i.e., volatile) and uninformative (i.e., stable) outcomes. Here, online prefrontal tDCS specifically enhanced learning rates when reward outcomes were *uninformative*. One explanation for this volatility-dependent effect of tDCS may be that there was greater scope for enhancing learning from these outcomes, as learning rates for *informative* wins were relatively high at baseline. In a replication study, however, we found no evidence for a volatility-dependent effect of tDCS on learning rates. Further work is therefore needed to determine the potential influence of volatility manipulations on tDCS outcomes.

Interestingly, DLPFC stimulation was associated with a reduction in participants’ total winnings. We hypothesize that this is related to the increased reliance on win outcomes independent of their true information content. It has previously been demonstrated that healthy adults are highly adept at modulating learning rates according to outcome volatility (Behrens et al., 2007; Nassar et al., 2012; Browning et al., 2015). Thus, if learning rates are (near-)optimal under normal conditions, stimulation-induced changes in either direction could cause poorer choices and thereby reduce winnings in this population. In individuals presenting with aberrant learning rates, on the other hand, prefrontal tDCS effects could potentially be harnessed to re-balance processing of positive and negative information.

## Limitations and future directions

This proof-of-concept study provides promising evidence for the use of prefrontal tDCS in ameliorating learning biases associated with mood disorders. However, there are several limitations which should be addressed in future studies. First, the replication study testing online prefrontal tDCS effects provided only a partial replication of the original findings. Specifically, there was no change in win-driven choice behaviour or total winnings. These differences may be explained as a relatively weaker effect of tDCS in the replication study, as non-model-based outcomes are less sensitive than the computational parameters. Additional independent replications would be useful to confirm the tDCS effects observed here. In addition, all tDCS effects were assessed in healthy adults with low scores on measures of depression or anxiety. Accumulating evidence shows that participants’ cognitive functioning, baseline electrophysiological state, and variation in recruitment of brain regions during task performance affect the outcomes of tDCS (Horvath et al., 2014; Li et al., 2015; Dubreuil-Vall et al., 2019). Compared to healthy adults, individuals with depression tend to be less sensitive to rewards (Eshel & Roiser, 2010) and present with hypoactivity of left DLPFC (Grimm et al., 2008; Disner et al., 2011). It can therefore not be ruled out that these neural and cognitive patterns would lead to different effects of prefrontal tDCS on learning performance. A logical next step within this line of research will be to determine whether the present findings can be extended to clinical groups. Finally, we did not examine transference of tDCS effects to different tasks or clinically meaningful outcomes. Further work is needed to address the potential generalisation of changes in reinforcement learning observed here.

## Conclusion

Using a computational approach, we demonstrated that reward learning rates can be modified in isolation with tDCS to bilateral DLPFC. tDCS effects were state-dependent, such that learning rates were increased with online but decreased with offline tDCS. These results provide preliminary evidence that online prefrontal tDCS can alter cognitive mechanisms critically associated with affective disorders. In the future, we aim to investigate whether this tDCS effect can be harnessed to shape learning processes and ameliorate clinical symptoms in individuals with depression.

## Acknowledgements

MJO was funded by a scholarship from the Medical Research Council. JO’S is a Sir Henry Dale Fellow funded by the Royal Society and the Wellcome Trust (HQR01720). The Wellcome Centre for Integrative Neuroimaging is supported by core funding from the Wellcome Trust (203139/Z/16/Z). VS was funded by a Medical Research Council studentship (MR/N013468/1). MB is supported by a Clinician Scientist Fellowship from the MRC (MR/N008103/1) and by the NIHR Oxford Health Biomedical Research Centre. M.J. Overman, V. Sarrazin and J. O’Shea declare that they have no conflict of interest. MB has received travel expenses from Lundbeck for attending conferences and acted as a consultant for Jansen Research and CHDR.

For the purpose of open access, the author has applied a CC BY public copyright license to any Author Accepted Manuscript version arising from this submission.

## Notes

https://osf.io/av9pf/

