## Supplemental Data for "Stimulating human prefrontal cortex increases reward learning"

***Extended Data***

***Model selection***

BIC values were calculated for the model used for analyses (**Model 1**) and the following five alternative models:

**Model 2:** Rather than modelling a single inverse temperature value, it is possible that participants display different stochastic choice behaviour for the win and loss outcomes. This model, which has been applied in previous studies of the IBLT (Pulcu & Browning, 2017), therefore incorporates separate inverse temperature parameters for wins and losses as follows:

$P_{(choice=A\left( i \right))}= \frac{1}{1+\exp^{(-{\beta win*rwin}_{\left( i \right)}-{\beta loss*rloss}_{\left( i \right)}))}}$	(1)

**Model 3:** Instead of learning the independent probabilities of win and loss outcomes, participants might take a model-free approach to the task by learning an overall value of each of the presented shapes (Pulcu & Browning, 2017):

${v^{A}}_{(i+1)}=v^{A}+ a*({out}_{\left( i \right)}- v_{(i)}^{A}$)	(2)

in which *v*^A^ represents the value of shape “A”, *α* is a single learning rate for updating the value, and *out*_(_*_i_*_)_ is the outcome of trial *i* (i.e., win – loss for shape “A”, which can be -1, 0, or 1). In the first trial, the value of shape “A” is set at 0. The estimated values of the two presented shapes were transformed into a choice probability by applying a softmax function using a single inverse temperature parameter.

**Model 4:** Similar to Model 1, this model calculates two learning rates and one inverse temperature parameter. This model is slightly simpler, however, in that it omits the “tendency” parameter *t* as described in Equation 4.

**Model 5:** A model using a single learning rate for win and loss outcomes combined, with two inverse temperature values for the two individual outcomes.

**Model 6:** The final model was similar to Model 2, with the main difference being that the values for the win and loss outcomes were centred at zero prior to multiplication with the inverse temperature values:

$P_{(choice=A\left( i \right))}= \frac{1}{1+\exp^{(-\left( {\beta win*(rwin}_{\left( i \right)}- 0.5 \right)-\left( {\beta loss*(rloss}_{\left( i \right)}- 0.5 \right))}}$	(3)

As shown in Fig.S.1, Model 1 provided the best fit to the data across the four studies and was therefore selected for the computational analyses.

Bayesian Information Criterion (BIC) values

|  | BIC |
| --- | --- |
| Model 1 | 547.626 |
| Model 2 | 550.968 |
| Model 3 | 617.937 |
| Model 4 | 556.589 |
| Model 5 | 555.148 |
| Model 6 | 566.1 |

**Figure S.1. Formal comparison of computational model fit.** Bars represent the sum of Bayesian Information Criterion (BIC) values per participant for each model. Smaller BIC values are indicative of a better model fit.

***Mood and anxiety measures***

To assess effects of tDCS on acute mood and anxiety, we contrasted participants’ questionnaire scores completed at the beginning and end of each session. Raw scores on each of the questionnaires are reported in Table S.1. Analyses were carried out using repeated-measures ANOVAs, with PANAS Positive scores, PANAS Negative scores, and STAI-State scores as dependent variables. tDCS and Time (before vs. after tDCS/task) were included as predictor variables. An interaction between tDCS and Time would be indicative of an effect of tDCS on acute mood or anxiety. We found no such interactions for PANAS Positive scores (*F*_(1,19)_ ≤ 3.68, *p* ≥ 0.072) or STAI-State scores (*F*_(1,19)_ _)_ ≤ 2.94, *p* ≥ 0.103). There was a significant effect of tDCS on PANAS Negative scores with offline prefrontal tDCS (tDCS x Time interaction: *F*_(1,19)_ = 5.72, *p* = 0.027, *η*^2^_G_ = 0.014). PANAS Negative scores decreased significantly with sham (*t*_(19)_ = 3.39, *p* = 0.003) but not active tDCS (*t*_(19)_ = 0.38, *p* = 0.705).

**Table S.1.** Mean (SD) PANAS and STAI-state scores by tDCS condition

|  | **Study 1**  *Online DLPFC  (N = 20)* | | **Study 2**  *Offline DLPFC*  *(N = 20)* | | **Study 3**  *Online M1*  *(N = 20)* | | **Study 4**  *Replication*  *(N = 20)* | |
| --- | --- | --- | --- | --- | --- | --- | --- | --- |
| ***PANAS Positive*** |  |  |  |  |  |  |  |  |
| Sham tDCS | 26.8 (4.9) | 22.4 (6.2) | 28.2 (6.4) | 26.0 (6.4) | 32.5 (7.3) | 30.4 (8.7) | 31.0 (7.10) | 27.4 (7.3) |
| Active tDCS | 27.2 (5.3) | 24.4 (6.6) | 28.6 (6.3) | 24.4 (6.0) | 31.3 (7.9) | 28.1 (8.6) | 30.6 (7.0) | 27.7 (8.6) |
| ***PANAS Negative*** |  |  |  |  |  |  |  |  |
| Sham tDCS | 10.9 (1.1) | 10.4 (0.6) | 13.0 (3.0) | 11.6 (2.6) | 11.0 (1.9) | 11.0 (1.4) | 11.5 (2.0) | 11.4 (2.4) |
| Active tDCS | 11.6 (2.3) | 10.6 (0.9) | 12.0 (2.7) | 11.9 (3.0) | 11.9 (4.6) | 10.8 (1.4) | 11.8 (2.1) | 11.1 (1.5) |
| ***STAI-State*** |  |  |  |  |  |  |  |  |
| Sham tDCS | 30.0 (4.7) | 31.8 (6.9) | 29.1 (6.7) | 30.0 (7.7) | 29.0 (7.3) | 29.8 (6.5) | 32.2 (8.7) | 30.6 (8.1) |
| Active tDCS | 30.0 (6.2) | 31.9 (4.8) | 30.5 (7.9) | 29.3 (6.7) | 30.6 (7.9) | 31.8 (6.9) | 29.6 (7.4) | 30.8 (8.6) |

*BDI-II; Beck’s Depression Inventory II, PANAS; Positive and Negative Affect Scale; STAI, State-Trait Anxiety Inventory.*

***Stability of prefrontal tDCS effects over time***
As shown in Figure S.2., the increase in reward learning rates in Win- and Loss-volatile blocks with online prefrontal tDCS was stable over time (tDCS x Outcome x Time interaction: *F*_(1,18)_ = 0.56, *p* = 0.464).

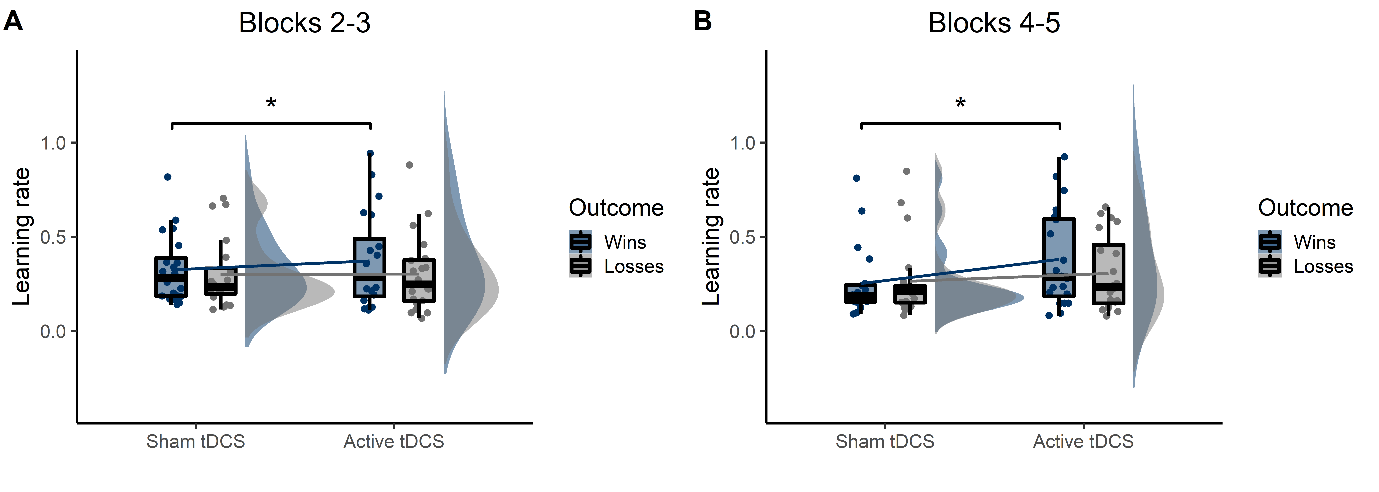

**Figure S.2. Stable increase in reward learning rates with online prefrontal tDCS (**p* <0.05). (A)** Learning rates in Blocks 2-3 (during tDCS). **(B)** Learning rates in Blocks 4-5 (after tDCS). Violin-plots show the distribution of learning rates by outcome (wins = blue, losses = grey). Summary statistics are provided in boxplots, with the black horizontal line indicating the median and whiskers representing the 25^th^ and 75^th^ percentiles of values. Dots represent participants’ individual data points averaged across task blocks.
